# DisCVR: Rapid viral diagnosis from high-throughput sequencing data

**DOI:** 10.1101/527127

**Authors:** Maha Maabar, Andrew J. Davison, Fiona Thorburn, Rory Gunson, Massimo Palmarini, Joseph Hughes

## Abstract

High-throughput sequencing (HTS) enables most pathogens in a clinical sample to be detected from a single analysis, thereby providing novel opportunities for diagnosis, surveillance and epidemiology. However, this powerful technology is difficult to apply in diagnostic laboratories because of its computational and bioinformatic demands. We have developed DisCVR, which detects known human viruses in clinical samples by matching sample *k*-mers (22 nucleotide sequences) to *k*-mers from taxonomically labelled viral genomes. DisCVR was validated using published HTS data for 89 clinical samples from adults with upper respiratory tract infections. These samples had been tested for viruses metagenomically and also by real-time polymerase chain reaction assay, which is the standard diagnostic method. DisCVR detected human viruses with high sensitivity (79%) and specificity (100%), and was able to detect mixed infections. Moreover, it produced results comparable to those in a published metagenomic analysis of 177 blood samples from patients in Nigeria. DisCVR has been designed as a user-friendly tool for detecting human viruses from HTS data using computers with limited RAM and processing power, and includes a graphical user interface to help users interpret and validate the output. It is written in Java and is publicly available from http://bioinformatics.cvr.ac.uk/discvr.php.

## 1. Introduction

The standard method for rapidly detecting known human viruses in clinical samples is the polymerase chain reaction (PCR), in which short oligonucleotides are used to amplify and probe specific regions of viral genomes. The limitations of this technique include the targeting of a relatively small number of viruses per assay and a dependence on sequence conservation among viral strains. High-throughput sequencing (HTS) provides approaches to viral diagnosis that have much greater scope. Thus, metagenomic analysis of HTS data can provide extensive viral genotyping information, as well as the characterization of complex multiple infections (Thorburn et al. 2015). Several metagenomic pipelines using *de novo* assembly and homology matching have been developed for virus detection (Li et al. 2016; Maarala et al. 2018; Ren et al. 2017; Scheuch et al. 2015; Wang et al. 2013; Zheng et al. 2017). However, analysing HTS data using such approaches brings heavy computing and bioinformatic demands that are difficult to meet and standardize in diagnostic laboratories (Flygare et al. 2016). As a consequence, we have developed DisCVR, which is a fast, accurate and easy-to-use tool for detecting known human viruses in clinical samples.

DisCVR employs an abundance-based method, which is a metagenomic approach for rapidly profiling the organisms present in a sample. It works by creating a database of short nucleotide sequences (*k*-mers) from a large set of viral reference sequences, tagging the *k*-mers taxonomically according to the viruses from which they came, screening each read in the HTS dataset for the presence of virus *k*-mers, and organising a summary of the viruses present in the sample via the tags. This approach makes data analysis very efficient, thereby minimizing the computing effort required (Orton et al. 2016).

Several existing tools utilize the abundance-based method to classify the reads in an HTS dataset. NBC (Rosen et al. 2011) employs a naïve Bayesian classifier to assign a log-likelihood score to each read. This classifier is trained by using a set of unique profiles of 15 nucleotide *k*-mers from microbial genomes, and then allows users to upload the dataset to a web site and obtain a summary of results listing the best taxonomic match for each read. Kraken (Wood & Salzberg 2014) assigns each *k*-mer in the database to the last common ancestor of species having that *k*-mer, and then assigns each read to the taxon with the most matching *k*-mers. CoMeta (Kawulok & Deorowicz 2015) creates a database of all *k*-mers for each rank in the taxonomic tree, and then uses these databases to classify the reads at each rank. CLARK (Ounit et al. 2015) collects target-specific *k*-mer sets from reference genomes belonging to a certain taxonomic rank (e.g. genus), and then classifies reads at that rank. This approach reduces the database size but requires a different database to be built for each rank. To improve the accuracy of the classification, CSSSCL (Borozan & Ferretti 2016) creates a BLAST database, a *k*-mer database and a compression database from a collection of reference genomes. Sequences in the sample are classified according to a combined sequence similarity score (CSSS) (Borozan et al. 2015) calculated from information in the pre-computed databases. In contrast to Kraken, CLARK and CoMeta, all of which assign individual reads, MetaPalette (Koslicki & Falush 2016) profiles the entire dataset and returns the relative proportions of organisms present by using *k*-mer sizes of 30 and 50, based on the rationale that using two different *k*-mer sizes allows strain-level variation to be captured more accurately. Taxonomer (Flygare et al. 2016) compares each read to multiple reference databases, assigning it to a high-level taxonomic category on the basis of the *k*-mer content of the read, and then uses exact *k*-mer matching to assign each read to a reference by maximizing the total *k*-mer weight. This weight, which is a function of the *k*-mer count in the reference and the database, provides a database-specific measure of how likely it is that a *k*-mer originated from a particular reference sequence.

Despite the growing number and popularity of *k*-mer-based classification tools, they have limitations. The databases are built using a limited set of reference sequences and therefore are of restricted utility for classifying organisms with sequences that diverge from the reference. This limitation can be a particular problem when significant variation exists in an organism at strain level. It can be addressed by incorporating a range of variants into the database, but this then creates a much larger database that may make the analysis challenging to run on resource-limited computers. Furthermore, many of the current tools are run on Linux systems and hence require the operator to have expertise in command line usage and an understanding of bioinformatics, which may be difficult to find in diagnostic settings. To our knowledge, the only tool that has been developed for ease of use and for application on computers with limited resources is Truffle (Visser et al. 2016). This is designed to screen for a limited set of user-specified viruses, comes preloaded with probe-sets for grapevine viruses, and cannot easily be updated for large sets of viruses from other hosts.

Here, we present DisCVR, a *k*-mer-based classification tool for detecting known human viruses from HTS data derived from clinical samples. DisCVR can be installed on a desktop computer to allow diagnostic laboratories to analyze large, confidential datasets by using a simple, straightforward graphical user interface (GUI) without specialized bioinformatics expertise. It is optimized to run on Windows, Linux and Mac OS, using minimal RAM and processing power without compromising speed and accuracy. The tool currently integrates curated viral databases at the taxonomic levels of species and strain, but may be used to build a customised database at any taxonomic level, thereby overcoming the limitations of using a restricted set of reference sequences. DisCVR utilizes *k*-mer counts derived from an entire HTS dataset to detect the viruses present in a sample, and validates the results by showing the coverage and depth of reads mapping to a reference sequence.

## 2. Methods

### 2.1 The *k*-mer databases

A *k*-mer is a short sequence of *k* nucleotides. A *k*-mer dataset is generated iteratively by sliding a window of size *k* along a sequence one nucleotide at a time. Extracting *k*-mers and counting their frequencies in a set of sequences can be computationally intensive, especially when *k* is large and the sequences are numerous. Dedicated *k*-mer counting programs, such as Jellyfish (Marçais & Kingsford 2011) and Khmer (Zhang et al. 2014), can be incorporated into abundance-based tools in order to optimize speed. KAnalyze (Audano & Vannberg 2014) was chosen for integration into DisCVR because the *k*-mers it generates are sorted lexicographically, thus making the search for matches very efficient. DisCVR also uses the canonical representation of a *k*-mer, which is lexicographically the smaller of a *k*-mer and its reverse complement.

For the purpose of this study, we define a virus *k*-mer as a *k*-mer that uniquely represents a virus or set of related viruses, to the exclusion of the host. A shared *k*-mer is defined as a *k*-mer that is common to a virus and the host. By excluding shared *k-*mers, it is not necessary for the user to remove host reads before using DisCVR, thus speeding up the overall processing time. If *k* is small, many copies of shared *k*-mers are generated, and if *k* is large, many copies of virus *k*-mers are found. Choosing the optimal *k*-mer size depends on balancing the advantages of speed (small *k*) with those of specificity and sensitivity (large *k*). Furthermore, it is necessary to reduce the number of low-complexity *k*-mers in the virus *k*-mer database, as these may be repetitive in sequence and present in otherwise unrelated viruses. The filtering of low-complexity *k*-mers and the selection of the size of *k* is explained in Supplementary Section S1.

For constructing the virus *k*-mer databases, three comprehensive datasets of complete or partial viral sequences were extracted from the NCBI taxonomy database. The first, the human hemorrhagic virus dataset (shortened below to “hemorrhagic dataset”), contained 33,367 sequences of the hemorrhagic fever viruses listed by the Centers for Disease Control and Prevention (‘Centers for Disease Control and Prevention’ n.d.). The second, the human respiratory virus dataset (“respiratory dataset”), contained 442,282 sequences of viruses associated with respiratory disease. The third, the human pathogenic virus dataset (“pathogenic dataset”), consisted of 1,762,968 sequences of viruses identified in the UK Health and Safety Executive list of biological agents (‘Health and Safety Executive: The approved list of biological agents.’ 2013).

### 2.2 Database build

DisCVR operates via three modules concerned with database build, sample classification and validation (Fig. 1).

**Figure 1.**
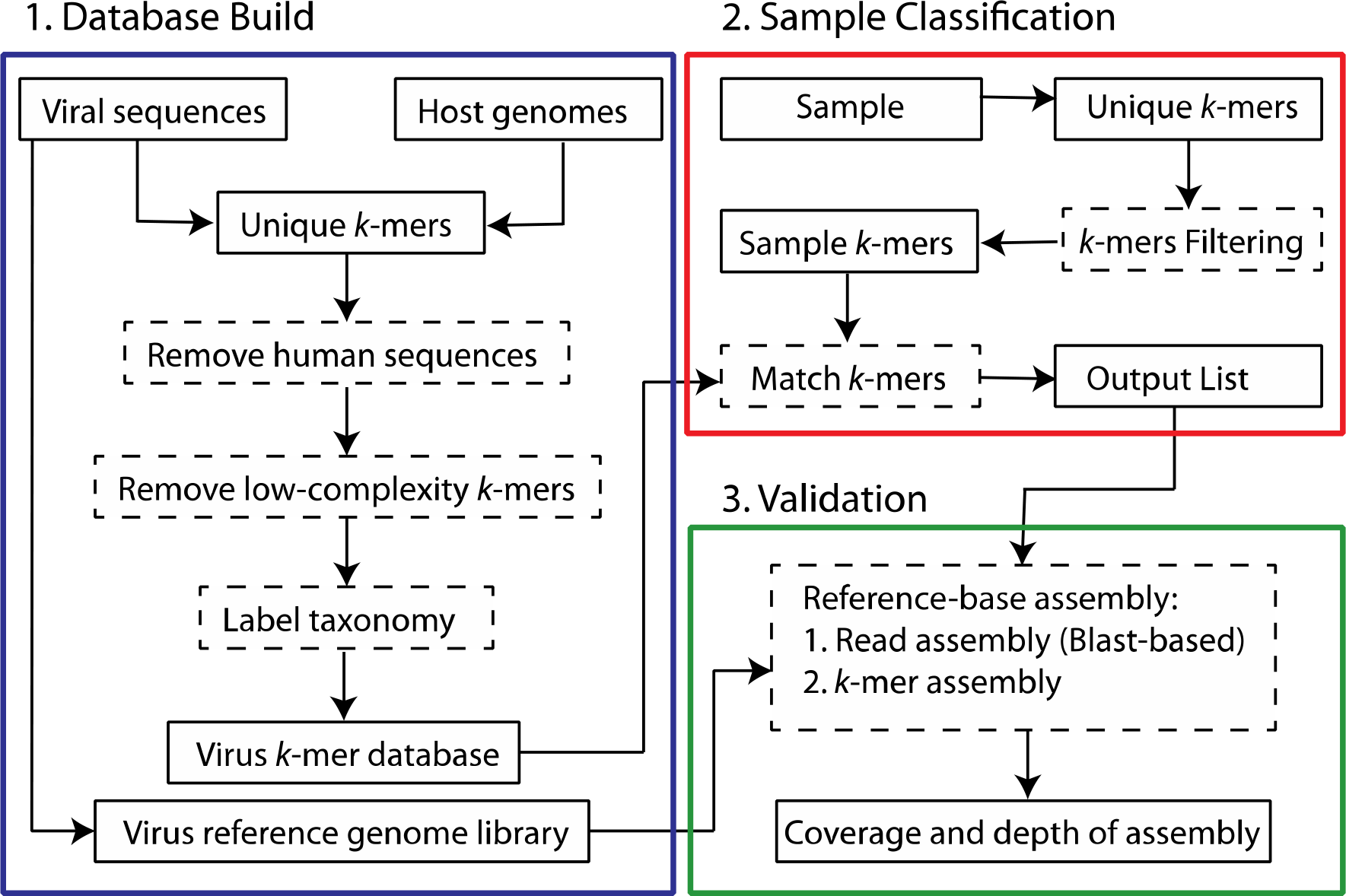
DisCVR framework. Each coloured box represents a component of the tool. Dashed rectangles indicate processes and solid rectangles show input and output.

Currently, the database build module includes three virus *k*-mer databases, derived from the hemorrhagic, respiratory and pathogenic datasets, for use in the sample classification module. In addition, some of the sequences in these datasets, defined largely by their presence in the NCBI RefSeq database, are used as a set of reference genome sequences in the validation module. DisCVR also allows the user to create customised databases and sets of reference sequences. The database build module involves selecting the relevant viral dataset, collecting the *k*-mers, and removing those that are shared with the host or are of low complexity. Each remaining *k*-mer is then identified with a taxonomic tag and an indication of the number of times it occurs in the sequences. The *k*-mers are further subdivided into those that exist in a single virus (i.e. specific *k*-mers) and those that exist in multiple viruses (i.e. non-specific *k*-mers). These assignments are made at the level of species and strain and are used in the output to illustrate the degree of specificity of the *k*-mers matching a virus (Fig. 2).

**Figure 2.**
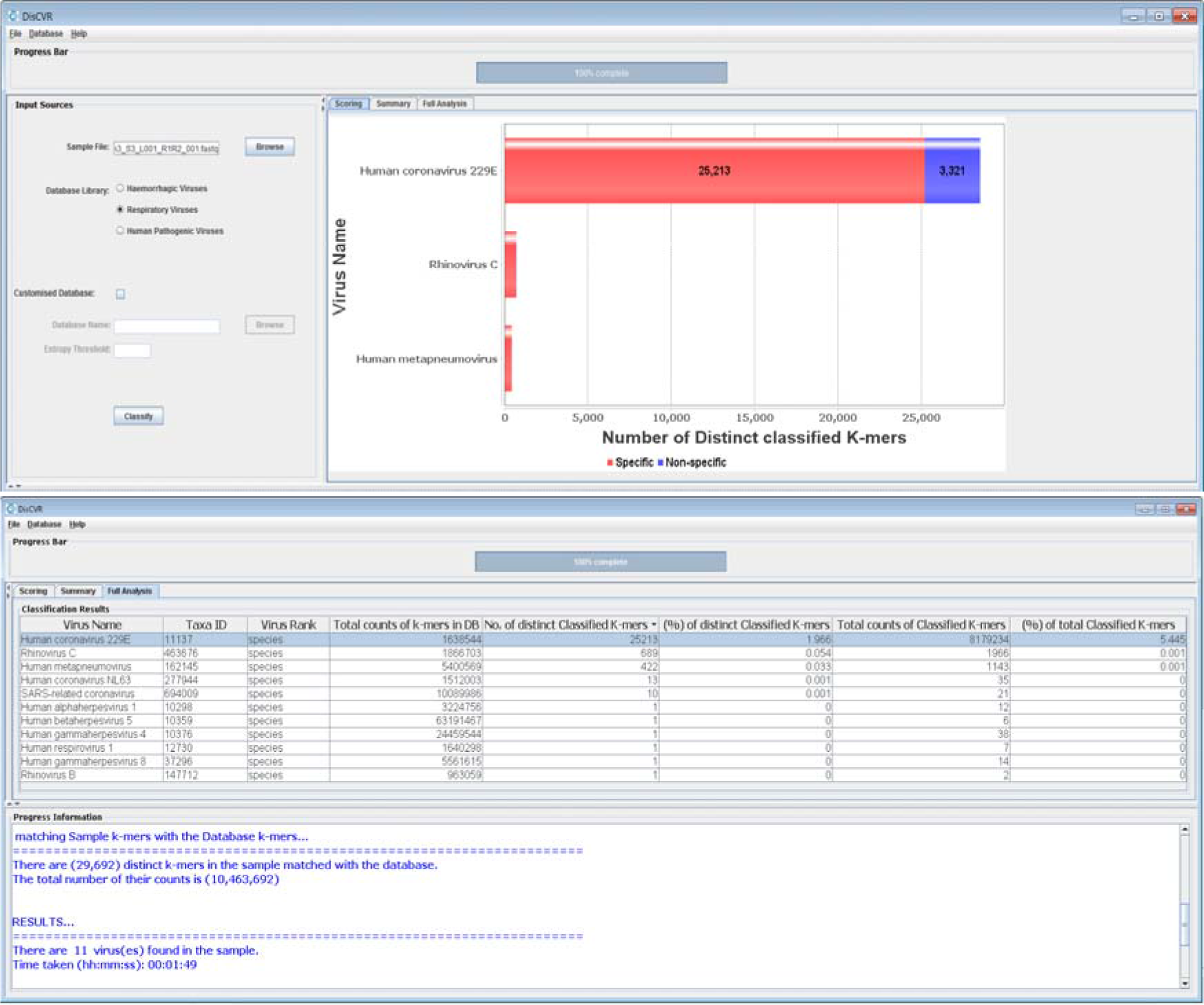
DisCVR GUI. The top screenshot shows the scoring panel with the top three virus hits, and the bottom screenshot shows the full analysis.

### 2.3 Sample classification

To analyze an HTS dataset, the file is loaded into DisCVR via the GUI. The *k*-mers are extracted and their frequencies are calculated, the single copy and low-complexity *k*-mers are filtered out, and the remaining *k*-mers are compared with the chosen virus *k*-mer database. As the number of *k*-mers in the sample can be enormous, various data structures were considered to optimize the classification on machines with limited RAM. Although searching the trie is fast O(n), where n is the size of the *k-*mer, it requires O(n^2^) overall time to build, and the space needed is quadratic. Instead, DisCVR uses a fast searching algorithm that groups similar *k*-mers together. Briefly, the *k*-mers in the virus database are divided among smaller sub-files according to the first five nucleotides. The same procedure is used to divide the *k*-mers derived from the entire HTS dataset. Searching commences by loading the corresponding sub-files from the virus *k*-mer database and the sample *k*-mers into memory, and performing a binary search for the presence of each sample *k*-mer among the database *k*-mers. Only matched *k*-mers are retrieved. Finally, DisCVR displays a straightforward list of all the virus hits detected, along with summary statistics and taxonomic information on the sample *k*-mers (Fig. 2).

### 2.4 Validation

DisCVR helps the user to assess the significance of the findings by facilitating an examination of *k*-mer distribution (allowing up to three mismatches) across a reference sequence representing the target genome. As an alternative, it also incorporates an examination of sequence read distribution carried out by using Tanoti (Sreenu n.d.), which is a BLAST-guided, reference-based short read aligner that is particularly tolerant of mismatches. In each case, the output is a graph showing the depth and coverage of *k*-mers or sequence reads across the reference genome and a summary of statistics for the mapping results (Fig. 3).

**Figure 3.**
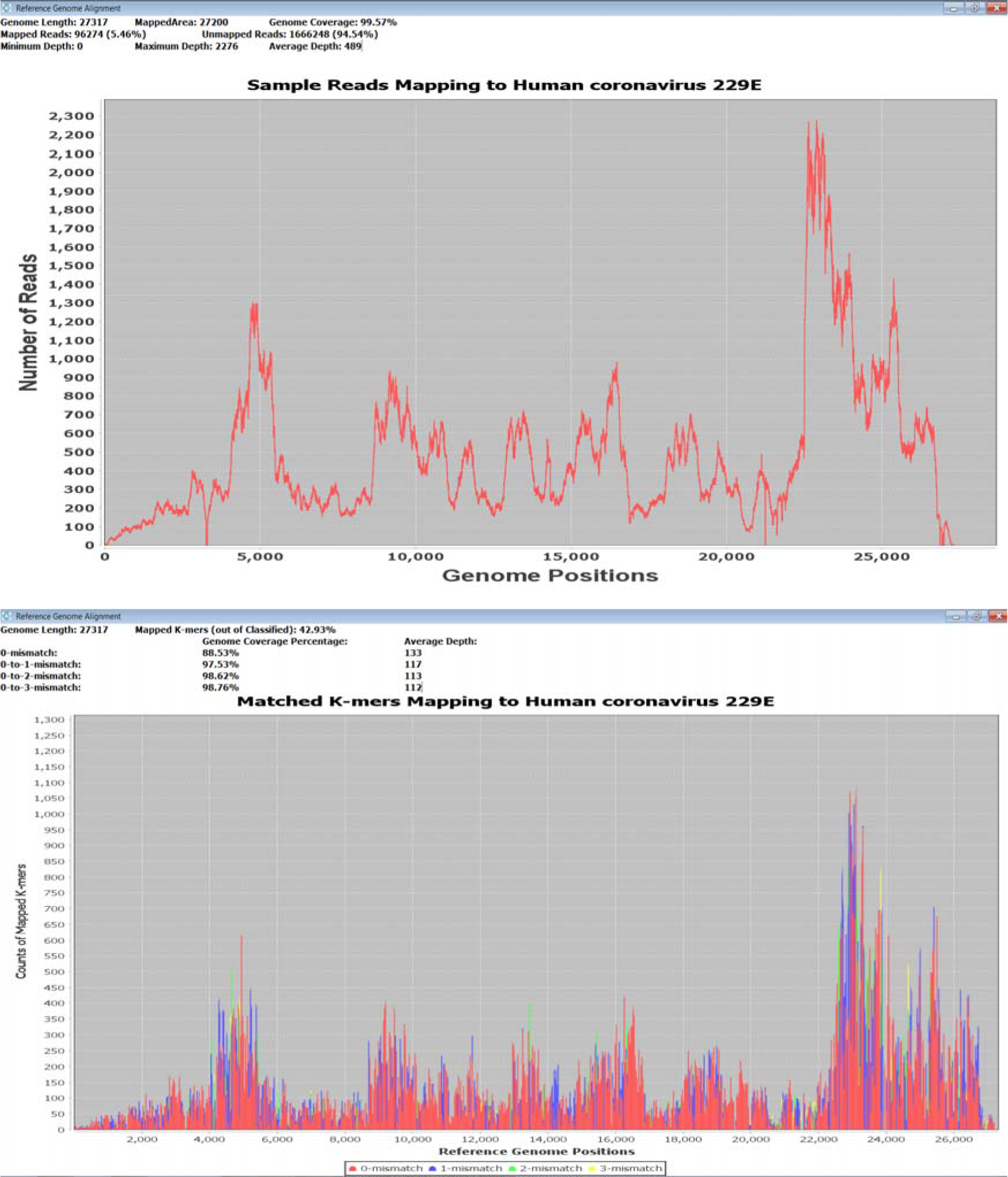
DisCVR validation. Coverage and depth of matched *k*-mers (top) and reads (bottom) to a reference genome.

### 2.5 Accuracy

The respiratory database was used to analyse published RNA-seq data from nasopharyngeal swab samples (n = 89) that had been collected from adults with upper respiratory tract infections (Thorburn et al. 2015) (Table S2; the average number of reads per sample was 660,640, range 30,872-1,278,122). The samples had been tested using a standard real-time PCR (RT-PCR) assay for human rhinovirus (HRV), influenza viruses A and B (IFA/IFB), respiratory syncytial virus (RSV), adenovirus (ADV), human metapneumovirus (hMPV), parainfluenza viruses (PIV) 1-4, and human coronaviruses (HCoV) HKU1, NL63, OC43 and 229E (Thorburn et al. 2015). The top hit for each sample (i.e. the virus having the greatest number of distinct *k*-mers) using DisCVR was compared to the virus detected previously by RT-PCR. The samples were also classified using three independent *k-*mer-based programs that require command-line usage on a Linux operating system: Kraken (Wood & Salzberg 2014), KrakenHLL (Breitwieser & Salzberg 2018) and CLARK (Ounit et al. 2015).

The initial objective was to determine the number of distinct *k*-mers that would maximize both sensitivity (effectiveness in identifying samples containing viruses) and specificity (effectiveness in identifying samples lacking viruses) for DisCVR. The output of DisCVR was categorized on the basis of the number of distinct *k*-mers for the top hit, and that of the other programs was assessed on the basis of the number of reads assigned to the top hit. For each tool, sensitivity and specificity were defined as TP/(TP+FN) and TN/(TN+FP), respectively, where TP, FN, TN and FP are the number of true positive, false negative, true negative and false positive samples relative to the RT-PCR results. We define samples as i) true positive when the top virus hit was detected by both RT-PCR and DisCVR, ii) true negative when neither RT-PCR nor DisCVR detected a virus, iii) false negative when a virus was detected by RT-PCR but not by DisCVR, and iii) false positive when a virus was detected by DisCVR but not by RT-PCR. ROC curves were generated for DisCVR, Kraken, KrakenHLL and CLARK using the pROC package in R and Youden’s statistic (Youden 1950).

### 2.6 Application

DisCVR was used to analyse 177 HTS RNA-seq libraries derived from serum specimens collected in Nigeria from healthy individuals (n = 120) and patients with unexplained acute febrile illness (n = 57) and analysed in a previous study (Stremlau et al. 2015). The raw data were downloaded from SRA BioProject PRJNA271229. The top hit using DisCVR was compared to the viral reads identified using BLASTn and BLASTx in the original study (https://doi.org/10.1371/journal.pntd.0003631.s017).

## 3. Results

The ROC curve (Fig. 4) derived from the datasets from respiratory tract infections (Thorburn et al. 2015) compares the sensitivity and specificity for different *k-*mer thresholds. It suggests that a value of 850 *k*-mers is the optimal threshold on the basis of the point on the curve furthest from the identity (diagonal) line (Table S2). The ROC curves of DisCVR and the other programs (Fig. 4) did not differ significantly from each other, and had overlapping confidence intervals. Kraken and KrakenHLL had identical curves. Kraken and CLARK rated as slightly more sensitive but less specific than DisCVR as a result of HCoV NL63 being the top hit in sample 1D3 and the second hit in DisCVR (Table 1 and Table S2). The top hit in DisCVR was HRV-A, which was the second hit in Kraken and CLARK but was not detected using RT-PCR. It was not informative to compare average execution time and memory usage for the programs, as it is not possible to run CLARK, Kraken and KrakenHLL natively on Windows operating systems.

**Figure 4.**
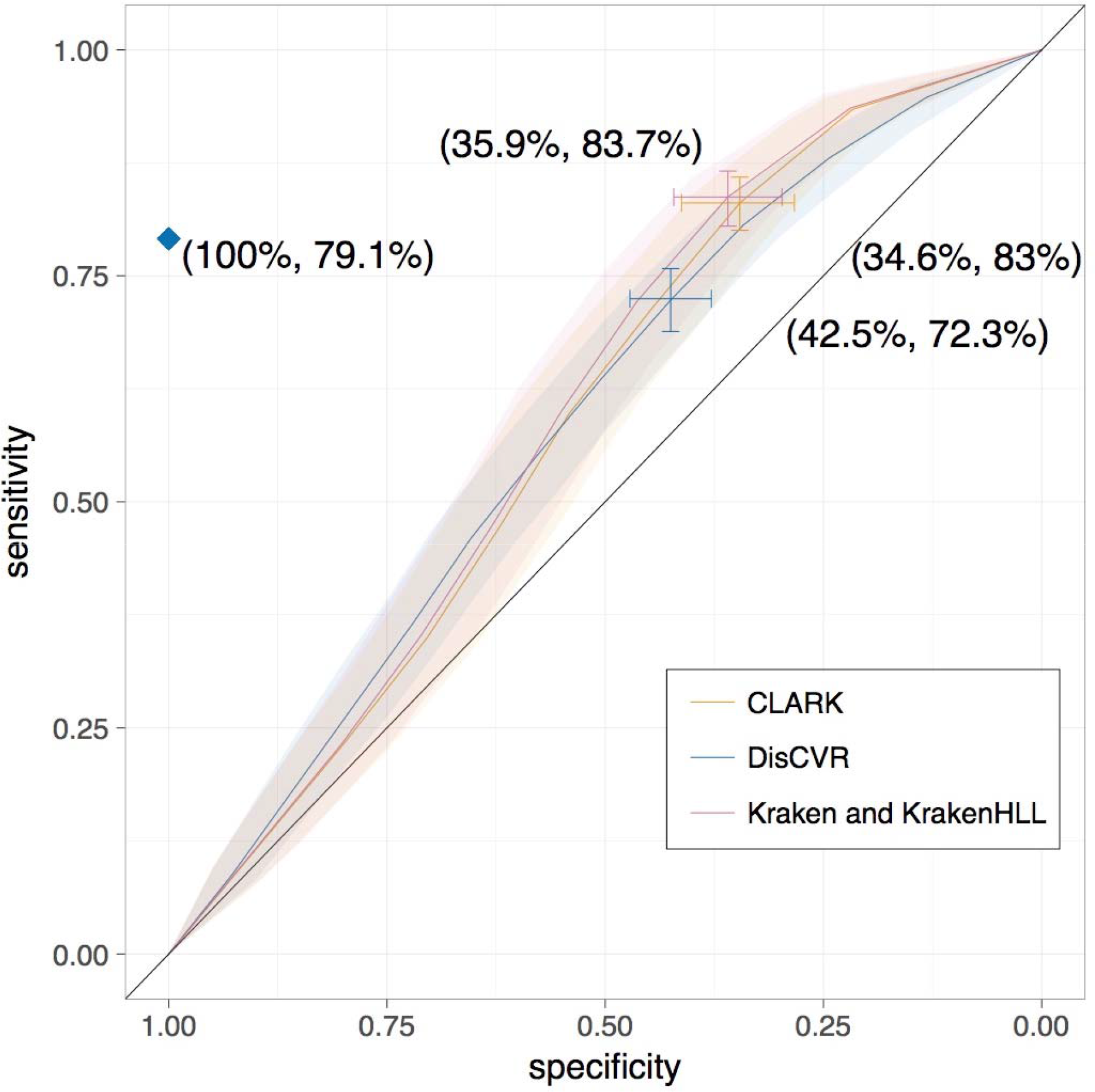
ROC curve showing the accuracy of DisCVR, CLARK and Kraken. The transparent shaded area shows the confidence interval of the sensitivity for all three methods. The optimal threshold of 850 *k-*mers for DisCVR and 150 reads for CLARK and Kraken are shown, with bars representing the confidence interval of the threshold and the specificity and sensitivity shown in brackets. The curve for KrakenHLL was identical to that for Kraken. The diamond indicates the sensitivity and specificity values, counting the false positives with ≥850 *k*-mers and the second hits with ≥850 *k*-mers among the true positives.

**Table 1.** Results of the second hits in the respiratory samples.

| Sample | RT-PCR<br>diagnosis | DisCVR<br>top hit and (no.)<sup>1</sup> | DisCVR<br>second hit and (no.)<sup>1</sup> |
| --- | --- | --- | --- |
| Top hit with ≤850 k-mers matching |  |  |  |
| 1G2 | PIV-3 | PIV-3 (366) | HRV-A (149) |
| 1I5 | HRV | HRV-A (749) | HRV-C (470) |
| 2B6 | RSV | RSV (742) | IFA H3N2 (262) |
| Second hit with ≥850 k-mers matching |  |  |  |
| 1B5 | PIV-3 | **HRV-A (3,758)** | **PIV-3 (3,111)** |
| 1D3 | HCoV NL63 | **HRV-A (2,420)** | **HCoV NL63 (1,841)** |
| Second hit with ≤850 k-mers matching |  |  |  |
| 1C2 | HRV | **Enterovirus D (1,633)** | HRV-A (269) |
| 1E5 | RSV | **HRV-C (1,777)** | RSV (415) |
| 1F8 | HCoV NL63 | **HRV-B (3,876)** | HCoV NL63 (724) |
| 2B9 | HRV | **RSV (1,105)** | HRV- C (94) |
| 2A2 | HCoV 229E | HRV-C (770) | HCoV 229E (176) |
| 2C4 | HCoV 229E | HRV-A (264) | HCoV 229E (5) |
| 2D3 | HCoV<br>OC43 | HRV-A (438) | HCoV OC43 (135) |
| 1F7 | HRV | hMPV (27) | HRV-B (20) |
| 1G1 | ADV/HRV | HCoV OC43 (163) | HRV-B (118) |
| Not detected |  |  |  |
| 1C9 | hMPV | **HRV-A (3,083)** | Enterovirus D (7) |
| 2D4 | PIV-2 | HRV-A (579) | HCoV OC43 (225) |

<sup>1</sup>Number of *k*-mers matching the classification. Hits with ≥850 k-mers are shown in bold.

A total of 48/89 (54%) of the samples had been shown to contain viruses by RT-PCR, and the remaining 41/89 lacked all viruses tested. Considering only the samples in the set of 89 for which DisCVR identified ≥850 *k*-mers for the top hit, the following findings were made. DisCVR identifed the viruses that were detected by RT-PCR in 32/48 (67%) of samples (true positives). It did not detect viruses in samples in which no viruses had been found by RT-PCR in 22/41 (54%) of samples (true negatives). It detected viruses in samples in which no viruses had been detected by RT-PCR in 19/41 (46%) of samples (false positives), and either detected viruses that did not correspond with those detected by RT-PCR or did not find any virus with ≥850 *k*-mers in 16/48 (33%) of samples (false negatives).

The RT-PCR assay was limited by the range of viruses that it could detect, by its dependence on sequence conservation, and consequently also by its potential to identify infections by multiple viruses. Consequently, the false positive results were assessed using the validation module (Table 2), and the false negative results were investigated by examining the second hits recorded by DisCVR (Table 1). In most false positive cases, the validation module showed that there were multiple reads mapping to several regions of the reference genome, thus confirming the presence of the viruses identified even though they had not been detected by RT-PCR. Some samples had low coverage because a single reference sequence (from RefSeq) represented the entire species but diverged in sequence from the virus present in the sample. For example, sample 1B3 yielded HRV-A89 (the reference for species *Rhinovirus A*) as the top hit, with only 7.6% genome coverage and 4 mapped reads. Using the capability of DisCVR to build a customized database drawn from the ≥100 prototypic strains of *Rhinovirus A*, HRV-A49 was revealed as the top hit, with 81.71% genome coverage and 263 mapped reads. This dramatic improvement illustrates the potential to strengthen the validation module by adding user-specific curated sets of sequences or by the proposed expansion of RefSeq entries capturing a greater degree of diversity (Brister et al. 2015). In the 16 false negative cases, DisCVR detected the virus identified by RT-PCR as the top hit in three samples (1G2, 1I5 and 2B6), but the number of distinct *k*-mers was <850 (Table 1 and Table S2). In addition, the virus identified by RT-PCR was detected as the second hit in 10 samples (1B5, 1D3, 1E5, 1G1, 1F7, 1F8, 2A2, 2B9, 2C4 and 2D3), and, in one case (1C2), the RT-PCR assay did not have the potential of identifying the top hit (enterovirus D). An important finding was made in two of these samples (1B5 and 1D3), in which the viruses detected by RT-PCR were not the top hits but still had ≥850 distinct *k*-mers in the sample (Table 1). This suggests that these patients were infected by multiple viruses. Finally, DisCVR did not detect any *k*-mers for the virus detected by RT-PCR in two samples (1C9 and 2D4), but identified HRV-A in 1C9, which was validated by reference assembly. The validation module thus yielded strong evidence for the presence of the viruses detected by DisCVR, at least where the number of *k*-mers was ≥850. These findings were taken into account in reassessing the sensitivity and specificity of DisCVR at 79% and 100%, respectively (Fig. 4).

**Table 2.** Coverage of reference genomes of the top hits detected in false positive samples in the respiratory samples.

| Sample | Virus detected<br>by DisCVR | Matched *k*-mers<sup>1</sup> | Genome coverage<br>(%) | No. mapped<br>reads (%)<sup>2</sup> |
| --- | --- | --- | --- | --- |
| 1B3 | HRV-A | 3,431 | 7.6 | 4 (0.00) |
| 1B4 | HRV-A | 3,652 | 9.39 | 14 (0.00) |
| 1B6 | HRV-A | 2,872 | 6.38 | 16 (0.00) |
| 1B9 | HRV-A | 1,041 | 2.15 | 1404 (0.10) |
| 1C8 | HRV-A | 2,781 | 8.21 | 8 (0.00) |
| 1D2 | HRV-A | 2,974 | 9.38 | 13 (0.00) |
| 1D5 | HRV-C | 901 | 3.63 | 8 (0.00) |
| 1D6 | HRV-C | 1,103 | 3.27 | 5 (0.99) |
| 1E2 | HRV-C | 1,299 | 1.51 | 1 (0.00) |
| 1E4 | HRV-C | 1,813 | 4.8 | 7 (0.00) |
| 1E9 | HRV-B | 4,306 | 13.69 | 27 (0.01) |
| 1G7 | HRV-B | 1,447 | 1.76 | 5 (0.00) |
| 1H5 | HRV-B | 932 | 3.84 | 4 (0.00) |
| 1I7 | HRV-C | 1,234 | 1.51 | 1 (0.00) |
| 1I9 | HRV-C | 1,845 | 3.1 | 9 (0.00) |
| 2A1 | RSV | 2,123 | 13.37 | 172 (0.02) |
| 2B5 | RSV | 927 | 13.56 | 69 (0.01) |
| 2B8 | RSV | 1,406 | 8.64 | 101 (0.01) |
| 2D1 | HRV-C | 1,620 | 1.59 | 2 (0.00) |

<sup>1</sup>Number of matching *k*-mers identified by the classification module.
<sup>2</sup>Percentage of total reads mapped by the validation module.

The threshold of 850 *k*-mers was also used in the analysis of the Nigerian datasets (Stremlau et al. 2015). The top hit from DisCVR was the same as that from the BLAST results in the original study for 101/177 (57%) cases, and viruses were detected in both healthy (n = 68) and afebrile (n = 33) patients (Table S4). In nine cases, the top hit from DisCVR differed from the top BLAST hit, but the second hit matched. In 55 cases, the number of *k-*mers was below the threshold in DisCVR, and the number of reads with BLAST matches was also low (an average of 24 reads per dataset). In the remaining 12 discordant samples, DisCVR detected human immunodeficiency virus 1 (n = 9), XMRV-related virus (n = 1) and human T-lymphotropic virus 1 (n = 1) as the top hit, whereas the BLAST results supported the presence of human adenovirus or Heterosigma akashiwo RNA virus (an algal virus). Mapping of reads to reference genomes suggested that the DisCVR and BLAST hits are false positives.

## 4. Discussion

Using HTS in diagnostic settings offers many advantages, including the ability to sequence pathogen genomes both individually and as communities. However, the uptake of HTS in such settings has been slow, due partly to the cost, turnover time and bioinformatic demands of this technology. We developed DisCVR to help address these challenges. DisCVR is a fast, accurate program for detecting viruses from HTS data using the increasingly exploited approach of *k*-mer classification. It offers the advantage of a non-targeted approach and also enables typing below the species level (e.g. subtype, serotype, genotype or strain). Unlike other tools for detecting viruses from HTS data, DisCVR is easy to use in diagnostic settings through the graphical user interface, requires no bioinformatic expertise, and can be used on the Windows operating systems that are commonly used in diagnostic laboratories. The basic output is easy to interpret, and the advanced output provides more detailed statistics and a validation capability.

DisCVR was designed for detecting known viruses and cannot be used to discover novel viruses. Indeed, the paper on the Nigerian patients (Stremlau et al. 2015) reported novel rhabdoviruses in healthy patients using a metagenomic approach, and these were not detected by DisCVR. However, metagenomics requires bioinformatic infrastructure and expertise at levels that are not commonly available in diagnostic laboratories. Nonetheless, DisCVR enables the detection of 148 pathogenic human viruses using one of the three implemented datasets (the pathogenic dataset), and more using the others. This represents a greater than ten-fold increase in target species over multiplex RT-PCR. Moreover, the number of viruses incorporated into the DisCVR databases is flexible, and can also be expanded by building custom databases.

In the datasets from respiratory tract infections, DisCVR had high sensitivity and specificity levels but did not identify all the viruses detected by RT-PCR when the threshold of ≥850 *k*-mers was used. This threshold may be set by the user and was calculated for the respiratory dataset for which we had paired RT-PCR and HTS data. As more datasets with paired information become available, it will be possible to tune the threshold more accurately to specific sample types and sizes. Further efforts could also be made to calibrate DisCVR from artificially constructed communities of viruses in various proportions.

Finally, DisCVR is configured as a human viral diagnostic tool, but could be readily expanded to include non-viral human pathogens and pathogens with non-human hosts by using the custom-build scripts in the DisCVR distribution.

## Supporting information

Supplementary Section S1

## Acknowledgements

We thank the members of the Viral Genomics and Bioinformatics Group for continuous, insightful feedback on DisCVR, and in particular Sejal Modha for generous support with utilizing NCBI tools and testing DisCVR. We also thank David Manlove for advice on algorithm design. This work was funded by the Medical Research Council (MC_UU_12014/12).

## Availability and requirements

Source code is available on github https://centre-for-virus-research.github.io/DisCVR/ and databases and executables are available on http://bioinformatics.cvr.ac.uk/discvr.php.

