## Supplementary Section S1 for "DisCVR: Rapid viral diagnosis from high-throughput sequencing data"

### Supplementary Methods for DisCVR: Rapid viral diagnosis from high-throughput sequencing data

#### Low-complexity *k*-mers

In order to exclude low-complexity *k*-mers, DisCVR calculates the tri-nucleotide Shannon entropy values of the *k*-mers (Shannon 1948). Examples consisting of a repeated nucleotide, a repeated tri-nucleotide and a sequence having no repeated pattern had entropy values of 0, 1 and ~3, respectively, thus demonstrating that this value is low for low-complexity *k*-mers (Supplementary Table S1). A program was written to extract *k*-mers of various sizes (*k*=18, 22, 26 or 30) from the pathogenic dataset, and their tri-nucleotide entropy values were calculated (Fig. S1). The distribution of entropy values showed that a threshold value of ≤2.5 excluded low-complexity *k*-mers while still including the majority of *k*-mers, with a larger proportion being retained as the size of *k* increased. This value was chosen as the default threshold setting built into DisCVR. The influence of filtering low-entropy *k*-mers is discussed below.

#### Optimal size of *k*

The proportions of virus and shared *k*-mers were calculated from the pathogenic dataset for increasing values of *k*, in relation to the total number of possible *k*-mers ($4^k$). The results show that the proportion of virus *k*-mers became greater than that of shared *k*-mers at *k*=18 (Fig. S2). This pattern was the same after eliminating low-complexity *k*-mers. An optimal *k*-mer size >18 was then chosen on the basis of processing time and number of virus *k*-mers.

The pathogenic and respiratory datasets were each used to construct virus *k*-mer databases at *k*=18, 22, 26 and 30 (Fig. S3). Each database was then used to analyse the published HTS data (Thorburn et al. 2015). The results confirmed that the smaller the *k*-mer, the less time taken, and the larger the *k*-mer, the lower the number of virus hits (Fig. S3). In addition, more time was required and more hits were obtained by using the pathogenic dataset, which is larger than the respiratory dataset.

Table S3 compares the results for the 48 samples that had tested positive by RT-PCR with the top scoring classification results obtained using the four databases generated from the respiratory dataset (*k*=18, 22, 26 and 30), with and without filtering low complexity *k*-mers (entropy ≤2.5). The observation that some of the classification results did not agree with the RT-PCR findings is discussed in the main text. Notably, the classification was insensitive to the size of *k*, with only two samples (1B5 and 1F7) not yielding the same result at all four sizes. Moreover, entropy filtering only influenced one sample at *k*=18 (1F7), where the top hit with filtering matched the RT-PCR assignment (HRV-B) but another target (human betaherpesvirus 5) without filtering (Table S2). Taking into account the results of these experiments, a default *k*-mer size of 22 was built into DisCVR. At this value, the average execution time per sample was 1 m 47 s on a Windows OS machine with 32 GB of RAM.

**Table S1**. Tri-nucleotide entropy values of repetitive and non-repetitive *k*-mers.

| *k* | *k*-mer sequence | Entropy |
|---|---|---|
| 18 | AAAAAAAAAAAAAAAAA | 0.0 |
| | TGTGTGTGTGTGTGTGTG | 1.0 |
| | GGTCCAGTGAAAGATCCT | 2.6 |
| 22 | AAAAAAAAAAAAAAAAAAAAAA | 0.0 |
| | TGTGTGTGTGTGTGTGTGTGTG | 1.0 |
| | GGTCCAGTGAAAGATCCTGTCA | 2.8 |
| 26 | AAAAAAAAAAAAAAAAAAAAAAAAA | 0.0 |
| | TGTGTGTGTGTGTGTGTGTGTGTGTG | 1.0 |
| | GGTCCAGTGAAAGATCCTGTCTATAG | 3.0 |
| 30 | AAAAAAAAAAAAAAAAAAAAAAAAAAAAA | 0.0 |
| | TGTGTGTGTGTGTGTGTGTGTGTGTGTGTG | 1.0 |
| | GGTCCAGTGAAAGATCCTGTCTATAGCATA | 3.3 |

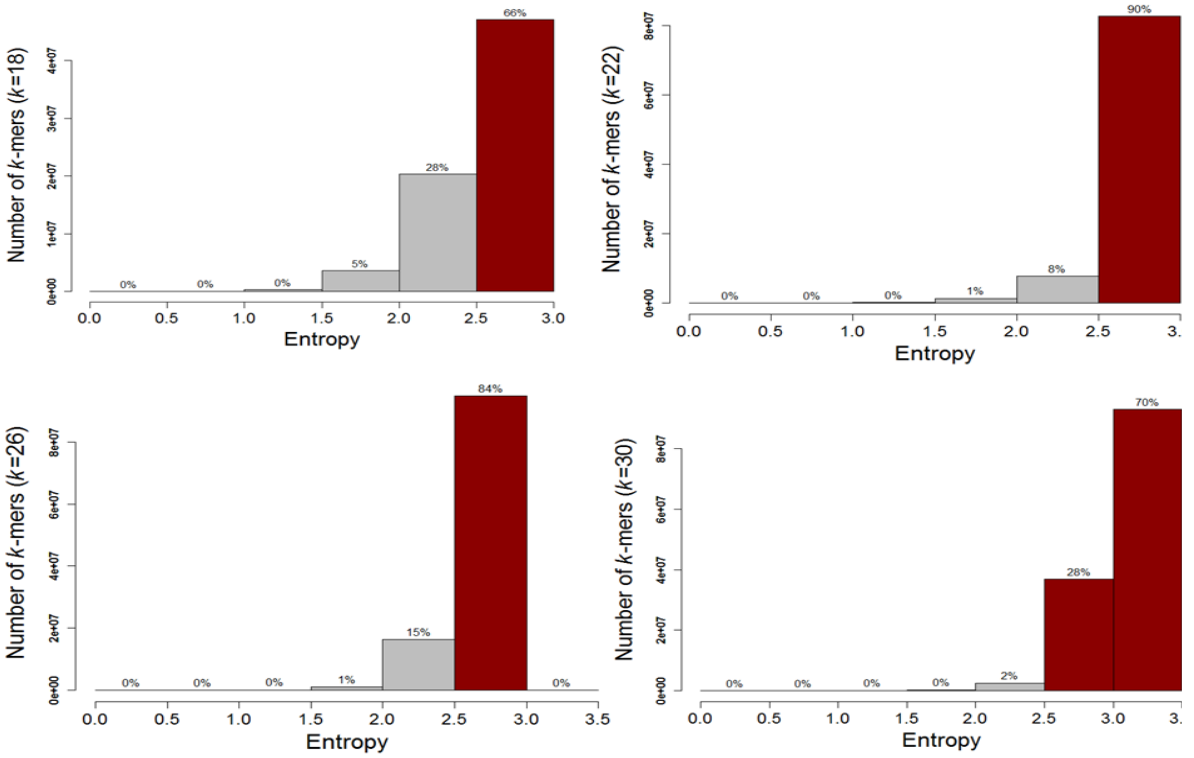


**Figure S1.** Entropy distribution for different values of *k*. *k*-mers above the threshold of 2.5 are indicated in red.

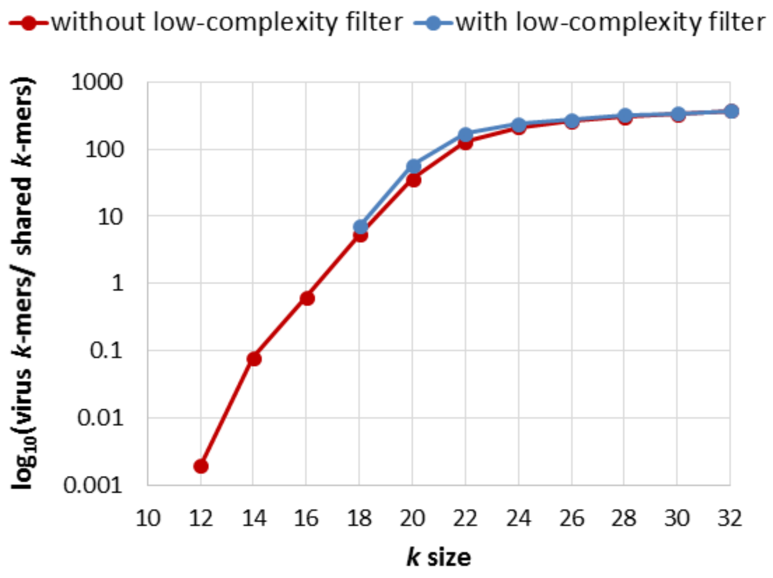


**Figure S2.** Relationship between the size of *k* and the proportion of virus *k*-mers in the pathogenic dataset. Low-complexity filter refers to the exclusion of low-complexity *k*-mers. All *k*-mers for *k*<18 were below the low-complexity threshold.

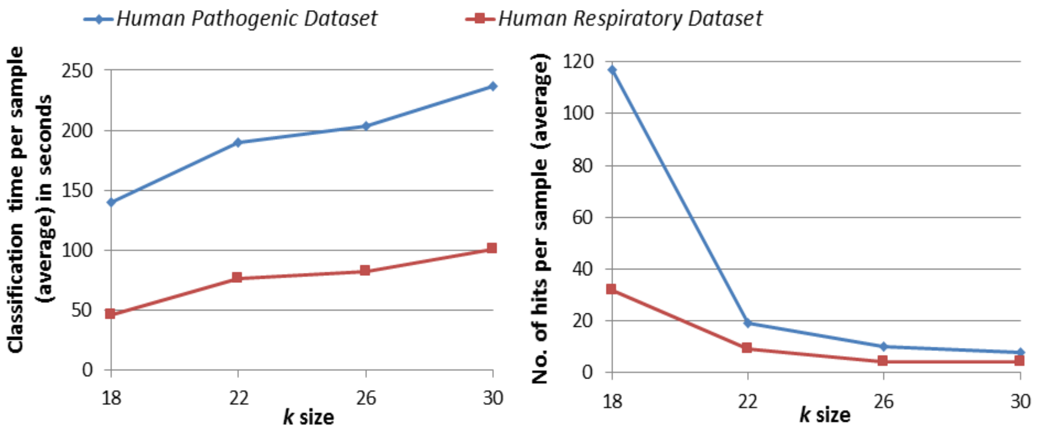


**Figure S3.** Relationship between *k*-mer size and classification results.
