## Supplementary Table S2 for "DisCVR: Rapid viral diagnosis from high-throughput sequencing data"

**Table S2: Comparison between RT-PCR results and DisCVR results in relation to the top two virus hits and the classification time.**

| Sample | # Reads | RT-PCR diagnosis | DisCVR first virus hit | Matched *k*-mers | DisCVR second virus hit | Matched *k*-mers | # Virus hits | Classification time (seconds) | CLARK top viral hit | No of matched reads with CLARK | Kraken top viral hit | No of matched reads with Kraken |
|---|---|---|---|---|---|---|---|---|---|---|---|---|
| 1B1 | 723,770 | HRV | HRV-A | 16,422 | HRV-C | 1,918 | 12 | 90 | Rhinovirus A | 538046 | Rhinovirus A | 529799 |
| 1B2 | 911,576 | HRV | HRV-A | 3,273 | HRV-B | 1,376 | 9 | 92 | Rhinovirus B | 19633 | Rhinovirus B | 11694 |
| 1B3 | 1,507,567 | | HRV-A | 3,431 | PIV-3 | 435 | 11 | 95 | Rhinovirus A | 436 | Rhinovirus A | 379 |
| 1B4 | 558,125 | | HRV-A | 3,652 | PIV-3 | 359 | 8 | 87 | Rhinovirus A | 466 | Rhinovirus A | 428 |
| 1B5 | 1,231,134 | PIV-3 | HRV-A | 3,758 | PIV-3 | 3,111 | 15 | 93 | Human respirovirus 3 | 59606 | Human respirovirus 3 | 59206 |
| 1B6 | 899,008 | | HRV-A | 2,872 | PIV-3 | 134 | 6 | 91 | Rhinovirus A | 263 | Rhinovirus A | 232 |
| 1B7 | 1,135,727 | HRV | HRV-A | 2,082 | HRV-C | 321 | 12 | 91 | Human gammaherpesvirus 4 | 13240 | Human gammaherpesvirus 4 | 36652 |
| 1B8 | 1,134,659 | IFA | IFA H3N2 | 1,043 | HRV-A | 50 | 8 | 90 | Influenza A virus | 336 | Influenza A virus | 333 |
| 1B9 | 1,397,408 | | HRV-A | 1,041 | PIV-3 | 21 | 12 | 96 | Rhinovirus A | 6785 | Rhinovirus A | 6516 |
| 1C2 | 2,195,230 | HRV | Enterovirus D | 1,633 | HRV-A | 269 | 12 | 101 | Enterovirus D | 21195 | Enterovirus D | 21108 |
| 1C3 | 2,248,306 | HRV | HRV-A | 2,585 | HRV-C | 1,827 | 13 | 101 | Rhinovirus B | 10630 | Rhinovirus B | 9566 |
| 1C4 | 473,341 | | HRV-C | 195 | HRV-A | 181 | 5 | 86 | Human alphaherpesvirus 1 | 26 | Human alphaherpesvirus 1 | 36 |
| 1C6 | 1,315,272 | | HRV-A | 90 | HRV-B | 23 | 7 | 91 | Human alphaherpesvirus 1 | 40 | Human alphaherpesvirus 1 | 30 |
| 1C7 | 835,392 | HRV | HRV-A | 4,289 | HRV-B | 11 | 7 | 87 | Rhinovirus A | 34741 | Rhinovirus A | 33871 |
| 1C8 | 1,154,678 | | HRV-A | 2,781 | HRV-C | 95 | 7 | 89 | Rhinovirus A | 339 | Rhinovirus A | 308 |
| 1C9 | 1,473,630 | hMPV | HRV-A | 3,083 | Enterovirus D | 7 | 10 | 94 | Rhinovirus A | 570 | Rhinovirus A | 523 |
| 1D1 | 1,426,921 | HRV | HRV-A | 11,900 | HRV-C | 163 | 12 | 94 | Rhinovirus A | 448384 | Rhinovirus A | 433462 |
| 1D2 | 1,762,196 | | HRV-A | 2,974 | HRV-C | 84 | 15 | 97 | Rhinovirus A | 420 | Rhinovirus A | 379 |
| 1D3 | 1,147,549 | HCoV NL63 | HRV-A | 2,420 | HCoV NL63 | 1,841 | 5 | 90 | Human coronavirus NL63 | 3141 | Human coronavirus NL63 | 3255 |
| 1D4 | 1,681,399 | HRV | HRV-A | 13,411 | HRV-C | 162 | 9 | 93 | Rhinovirus A | 1311180 | Rhinovirus A | 1247916 |
| 1D5 | 1,568,303 | | HRV-C | 901 | HRV-A | 11 | 6 | 93 | Rhinovirus C | 73 | Human alphaherpesvirus 1 | 83 |
| 1D6 | 1,208,371 | | HRV-C | 1,103 | HRV-B | 547 | 14 | 92 | Rhinovirus B | 165 | Rhinovirus B | 157 |
| 1D7 | 2,120,505 | | HRV-A | 581 | HCoV OC43 | 117 | 14 | 96 | Rhinovirus A | 122 | Human alphaherpesvirus 1 | 79 |
| 1D8 | 1,285,318 | | HRV-A | 466 | PIV-3 | 194 | 12 | 91 | Rhinovirus A | 100 | Human alphaherpesvirus 1 | 68 |
| 1D9 | 1,636,891 | | HRV-A | 268 | PIV-3 | 20 | 13 | 94 | Rhinovirus A | 89 | Rhinovirus A | 52 |
| 1E1 | 1,038,182 | HRV | HRV-C | 11,268 | HRV-A | 1,311 | 14 | 92 | Rhinovirus C | 321843 | Rhinovirus C | 313519 |
| 1E2 | 685,534 | | HRV-C | 1,299 | RSV | 288 | 6 | 92 | Rhinovirus C | 66 | Rhinovirus C | 64 |
| 1E3 | 919,930 | RSV | RSV | 19,820 | HRV-C | 1,471 | 11 | 92 | Human orthopneumovirus | 861892 | Human orthopneumovirus | 30061 |
| 1E4 | 803,117 | | HRV-C | 1,813 | RSV | 360 | 8 | 91 | Rhinovirus C | 134 | Rhinovirus C | 129 |
| 1E5 | 956,806 | RSV | HRV-C | 1,777 | RSV | 415 | 9 | 94 | Rhinovirus C | 103 | Rhinovirus C | 90 |
| 1E6 | 278,332 | | HRV-B | 181 | RSV | 39 | 2 | 85 | Human alphaherpesvirus 1 | 69 | Human alphaherpesvirus 1 | 132 |
| 1E7 | 777,070 | HCoV NL63 | HCoV NL63 | 5,075 | HRV-B | 22 | 8 | 91 | Human coronavirus NL63 | 4430 | Human coronavirus NL63 | 4414 |
| 1E8 | 1,330,431 | HRV | HRV-C | 1,384 | HCoV 229E | 115 | 8 | 97 | Rhinovirus C | 574 | Rhinovirus C | 559 |
| 1E9 | 207,805 | | HRV-B | 4,306 | RSV | 32 | 4 | 84 | Rhinovirus B | 659 | Rhinovirus B | 577 |
| 1F1 | 685,468 | RSV | RSV | 7,502 | HCoV OC43 | 84 | 5 | 86 | Human orthopneumovirus | 13475 | Human orthopneumovirus | 13413 |
| 1F2 | 729,478 | | HRV-A | 211 | RSV | 94 | 7 | 85 | Human alphaherpesvirus 1 | 52 | Human alphaherpesvirus 1 | 99 |
| 1F3 | 936,244 | | HRV-B | 52 | HHV-5 | 1 | 4 | 87 | Human alphaherpesvirus 1 | 89 | Human alphaherpesvirus 1 | 171 |
| 1F5 | 1,012,230 | | HRV-B | 39 | HHV-4 | 1 | 6 | 88 | Human alphaherpesvirus 1 | 104 | Human alphaherpesvirus 1 | 198 |
| 1F7 | 798,626 | HRV | hMPV | 27 | RSV | 21 | 6 | 86 | Human alphaherpesvirus 1 | 90 | Human alphaherpesvirus 1 | 204 |
| 1F8 | 1,347,125 | HCoV NL63 | HRV-B | 3,876 | HCoV NL63 | 724 | 6 | 90 | Rhinovirus B | 343 | Rhinovirus B | 292 |
| 1G1 | 831,318 | HRV/ADV | HCoV OC43 | 163 | HRV-B | 118 | 7 | 86 | Rhinovirus A | 90 | Human alphaherpesvirus 1 | 98 |
| 1G2 | 1,077,226 | PIV-3 | PIV-3 | 366 | HRV-A | 149 | 7 | 86 | Human respirovirus 3 | 122 | Human alphaherpesvirus 1 | 144 |
| 1G5 | 751,698 | | HRV-A | 64 | HHV-4 | 3 | 8 | 89 | Human alphaherpesvirus 1 | 75 | Human alphaherpesvirus 1 | 92 |
| 1G6 | 355,688 | hMPV | hMPV | 1,945 | HCoV OC43 | 127 | 5 | 82 | Human metapneumovirus | 1355 | Human metapneumovirus | 1324 |
| 1G7 | 979,331 | | HRV-B | 1,447 | HRV-A | 40 | 6 | 87 | Rhinovirus B | 104 | Rhinovirus B | 84 |
| 1H1 | 827,978 | HCoV OC43 | HCoV OC43 | 27,483 | RSV | 89 | 11 | 90 | Betacoronavirus 1 | 23000 | Human coronavirus OC43 | 22891 |

| | | | | | | | | | | | | |
|---|---|---|---|---|---|---|---|---|---|---|---|---|
| 1H3 | 735,513 | HRV | HRV-A | 3,390 | HRV-B | 36 | 6 | 86 | Rhinovirus A | 51859 | Rhinovirus A | 39470 |
| 1H4 | 681,586 | | HRV-B | 431 | RSV | 2 | 3 | 84 | Human alphaherpes virus 1 | 65 | Human alphaherpes virus 1 | 135 |
| 1H5 | 1,515,119 | | HRV-B | 932 | HHV- 4 | 3 | 7 | 94 | Human alphaherpes virus 1 | 96 | Human alphaherpes virus 1 | 201 |
| 1H6 | 671,898 | | HRV-B | 696 | RSV | 19 | 4 | 85 | Human alphaherpes virus 1 | 75 | Human alphaherpes virus 1 | 135 |
| 1H7 | 1,167,844 | HRV | HRV-B | 8,718 | HRV-A | 116 | 7 | 89 | Rhinovirus B | 330281 | Rhinovirus B | 324189 |
| 1H8 | 767,616 | | HRV-C | 811 | HRV-A | 363 | 12 | 86 | Rhinovirus C | 79 | Rhinovirus C | 54 |
| 1I2 | 967,819 | HRV | HRV-C | 8,356 | HRV-A | 533 | 7 | 91 | Rhinovirus C | 312810 | Rhinovirus C | 303380 |
| 1I3 | 1,101,849 | | HRV-C | 695 | HRV-A | 14 | 13 | 91 | Rhinovirus C | 46 | Rhinovirus C | 37 |
| 1I4 | 875,020 | HRV | HRV-B | 1,011 | HRV-C | 812 | 9 | 90 | Rhinovirus B | 2838 | Rhinovirus B | 2237 |
| 1I5 | 1,090,702 | HRV | HRV-A | 749 | HRV-C | 470 | 4 | 89 | Rhinovirus A | 107 | Rhinovirus A | 75 |
| 1I6 | 1,636,747 | HRV | HRV-C | 7,801 | HRV-A | 240 | 7 | 97 | Rhinovirus C | 499178 | Rhinovirus C | 410983 |
| 1I7 | 1,357,899 | | HRV-C | 1,243 | HHV-4 | 6 | 7 | 93 | Rhinovirus C | 105 | Rhinovirus C | 78 |
| 1I8 | 2,149,357 | HRV | HRV-C | 1,026 | HRV-B | 282 | 7 | 97 | Rhinovirus B | 188 | Rhinovirus B | 171 |
| 1I9 | 1,376,809 | | HRV-C | 1,845 | HRV-A | 40 | 3 | 89 | Rhinovirus C | 169 | Rhinovirus C | 142 |
| 2A1 | 808,953 | | RSV | 2,123 | HRV-A | 1173 | 7 | 88 | Human orthopneum ovirus | 328 | Human orthopneum ovirus | 282 |
| 2A2 | 1,359,030 | HCoV 229E | HRV-C | 770 | HCoV 229E | 176 | 7 | 92 | Rhinovirus C | 67 | Rhinovirus C | 56 |
| 2A3 | 1,771,886 | HCoV 229E | HCoV 229E | 25,213 | HRV-C | 689 | 11 | 95 | Human coronavirus 229E | 114203 | Human coronavirus 229E | 113611 |
| 2A4 | 496,185 | HCoV 229E | HCoV 229E | 7,515 | HRV-C | 477 | 9 | 64 | Human coronavirus 229E | 512 | Human coronavirus 229E | 512 |
| 2A5 | 1,172,473 | | HRV-C | 486 | HCoV 229E | 141 | 6 | 68 | Rhinovirus C | 40 | Rhinovirus C | 37 |
| 2A6 | 1,447,894 | HRV | HRV-C | 6,136 | HRV-A | 160 | 13 | 93 | Rhinovirus C | 150810 | Rhinovirus C | 143621 |
| 2A7 | 961,849 | | RSV | 121 | hMPV | 69 | 5 | 85 | Human orthopneum ovirus | 16 | Human orthopneum ovirus | 13 |
| 2A8 | 957,931 | | hMPV | 29 | HHV-4 | 3 | 6 | 86 | Rhinovirus C | 9 | Human alphaherpes virus 1 | 13 |
| 2A9 | 1,354,113 | hMPV | hMPV | 8,428 | HCoV 229E | 41 | 12 | 90 | Human metapneum ovirus | 50724 | Human metapneum ovirus | 50085 |
| 2B1 | 1,177,323 | | IFA H3N2 | 101 | IFA H1N1 | 47 | 10 | 89 | Influenza A virus | 531 | H3N2 subtype | 344 |
| 2B2 | 1383897 | | hMPV | 48 | HRV-A | 16 | 8 | 89 | Human alphaherpes virus 1 | 18 | Human alphaherpes virus 1 | 22 |
| 2B3 | 1,172,718 | | HRV-C | 33 | hMPV | 10 | 10 | 89 | Rhinovirus C | 24 | Human alphaherpes virus 1 | 23 |
| 2B4 | 1,217,147 | RSV | RSV | 40,911 | HRV-A | 65 | 8 | 87 | Human orthopneum ovirus | 343702 | Human orthopneum ovirus | 339723 |
| 2B5 | 1,029,140 | | RSV | 927 | HRV-C | 3 | 4 | 88 | Human orthopneum ovirus | 150 | Human orthopneum ovirus | 137 |
| 2B6 | 1,214,549 | RSV | RSV | 742 | IFA H3N2 | 262 | 14 | 90 | Influenza A virus | 324 | H3N2 subtype | 184 |
| 2B7 | 925,821 | HRV | HRV-A | 4,458 | RSV | 1,553 | 6 | 86 | Rhinovirus A | 13289 | Rhinovirus A | 12362 |
| 2B8 | 886,038 | | RSV | 1,406 | HRV-C | 5 | 6 | 87 | Human orthopneum ovirus | 169 | Human orthopneum ovirus | 158 |
| 2B9 | 896,407 | HRV | RSV | 1,105 | HRV-C | 94 | 3 | 85 | Human orthopneum ovirus | 141 | Human orthopneum ovirus | 133 |
| 2C1 | 1,521,785 | HRV | HRV-A | 7631 | HRV-C | 46 | 9 | 92 | Rhinovirus A | 50040 | Rhinovirus A | 48395 |
| 2C2 | 1,328,935 | | HRV-A | 42 | HRV-C | 4 | 10 | 92 | Rhinovirus C | 23 | Rhinovirus C | 9 |
| 2C3 | 811,298 | HCoV 229E | HCoV 229E | 3169 | HRV-A | 270 | 6 | 88 | Human coronavirus 229E | 2411 | Human coronavirus 229E | 2380 |
| 2C4 | 1,152,141 | HCoV 229E | HRV-A | 264 | HCoV 229E | 5 | 7 | 89 | Rhinovirus A | 11 | Rhinovirus A | 11 |
| 2C6 | 1,379,271 | | HRV-A | 57 | HRV-C | 4 | 8 | 92 | Rhinovirus C | 16 | Human alphaherpes virus 1 | 25 |
| 2D1 | 1,145,000 | | HRV-C | 1,620 | RSV | 215 | 9 | 87 | Rhinovirus C | 91 | Rhinovirus C | 86 |
| 2D2 | 1,008,557 | HCoV 229E | HCoV 229E | 14,279 | HRV-A | 795 | 11 | 88 | Human coronavirus 229E | 20065 | Human coronavirus 229E | 19979 |
| 2D3 | 1,223,798 | HCoV OC43 | HRV-A | 438 | HCoV OC43 | 135 | 10 | 87 | Rhinovirus A | 124 | Human alphaherpes virus 2 | 221 |
| 2D4 | 1,473,622 | PIV-2 | HRV-A | 579 | HCoV OC43 | 225 | 12 | 94 | Rhinovirus A | 156 | Rhinovirus A | 90 |
| 2D5 | 1,381,472 | HCoV OC43 | HCoV OC43 | 26,870 | HRV-A | 318 | 17 | 91 | Betacoronav irus 1 | 30089 | Human coronavirus OC43 | 29799 |
| 2D6 | 1,374,102 | HRV | HRV-A | 3,700 | HCoV OC43 | 238 | 11 | 91 | Rhinovirus A | 100574 | Rhinovirus A | 53062 |
