## Supplementary Table S3 for "DisCVR: Rapid viral diagnosis from high-throughput sequencing data"

**Table S3: Comparison between published RT-PCR results and DisCVR results in relation to *k*-mer size.**

| Sample | RT-PCR detection | *k* = 18 | Matched k-mers | *k* = 22 | Matched *k*-mers | *k* = 26 | Matched *k*-mers | *k* = 30 | Matched *k*-mers |
|---|---|---|---|---|---|---|---|---|---|
| 1B1 | HRV | HRV-A | 12,385 | HRV-A | 16,422 | HRV-A | 13,424 | HRV-A | 13,730 |
| 1B2 | HRV | HRV-A | 2,190 | HRV-A | 3,273 | HRV-A | 2,827 | HRV-A | 2,935 |
| 1B5 | PIV-3 | HRV-A | 2,635 | HRV-A | 3,758 | HRV-A | 3,219 | PIV-3 | 3,951 |
| 1B7 | HRV | HRV-A | 1,572 | HRV-A | 2,082 | HRV-A | 1,593 | HRV-A | 1,531 |
| 1B8 | IFA | IFA H3N2 | 637 | IFA H3N2 | 1,043 | IFA H3N2 | 1,029 | IFA H3N2 | 1,225 |
| 1C2 | HRV | Enterovirus D | 1,056 | Enterovirus D | 1,633 | Enterovirus D | 1,524 | Enterovirus D | 1,686 |
| 1C3 | HRV | HRV-A | 1,766 | HRV-A | 2,585 | HRV-A | 2,220 | HRV-A | 2,238 |
| 1C7 | HRV | HRV-A | 2,876 | HRV-A | 4,289 | HRV-A | 3,707 | HRV-A | 3,828 |
| 1C9 | hMPV | HRV-A | 2,081 | HRV-A | 3,083 | HRV-A | 2,695 | HRV-A | 2,779 |
| 1D1 | HRV | HRV-A | 9,283 | HRV-A | 11,900 | HRV-A | 9,226 | HRV-A | 8,813 |
| 1D3 | HCoV NL63 | HRV-A | 1,670 | HRV-A | 2,420 | HRV-A | 2,107 | HRV-A | 2,120 |
| 1D4 | HRV | HRV-A | 10,589 | HRV-A | 13,411 | HRV-A | 10,112 | HRV-A | 9,445 |
| 1E1 | HRV | HRV-C | 8,349 | HRV-C | 11,268 | HRV-C | 10,046 | HRV-C | 10,428 |
| 1E3 | RSV | RSV | 11,002 | RSV | 19,820 | RSV | 18,013 | RSV | 22,161 |
| 1E5 | RSV | HRV-C | 1,315 | HRV-C | 1,777 | HRV-C | 1,454 | HRV-C | 1,447 |
| 1E7 | HCoV NL63 | HCoV NL63 | 2,881 | HCoV NL63 | 5,075 | HCoV NL63 | 4,702 | HCoV NL63 | 5,511 |
| 1E8 | HRV | HRV-C | 989 | HRV-C | 1,384 | HRV-C | 1,289 | HRV-C | 1,381 |
| 1F1 | RSV | RSV | 4,265 | RSV | 7,502 | RSV | 6,945 | RSV | 8,018 |
| 1F7 | HRV | HRV-B | 31 | hMPV | 27 | hMPV | 23 | hMPV | 19 |
| 1F8 | HCoV NL63 | HRV-B | 2,624 | HRV-B | 3,876 | HRV-B | 3,414 | HRV-B | 3,545 |
| 1G1 | HRV/ADV | HCoV OC43 | 109 | HCoV OC43 | 163 | HCov OC43 | 129 | HCoV OC43 | 147 |
| 1G2 | PIV-3 | PIV-3 | 233 | PIV-3 | 366 | PIV-3 | 333 | PIV-3 | 387 |
| 1G6 | hMPV | hMPV | 1,126 | hMPV | 1,945 | hMPV | 1,751 | hMPV | 2,004 |
| 1H1 | HCoV OC43 | HCoV OC43 | 17,031 | HCoV OC43 | 27,483 | HCov OC43 | 25,792 | HCoV OC43 | 29,918 |
| 1H3 | HRV | HRV-A | 2,813 | HRV-A | 3,390 | HRV-A | 2,471 | HRV-A | 2,339 |
| 1H7 | HRV | HRV-B | 6,487 | HRV-B | 8,718 | HRV-B | 7,349 | HRV-B | 7,414 |
| 1I2 | HRV | HRV-C | 6,556 | HRV-C | 8,356 | HRV-C | 7,110 | HRV-C | 7,123 |
| 1I4 | HRV | HRV-B | 657 | HRV-B | 1,011 | HRV-B | 943 | HRV-B | 1,074 |
| 1I5 | HRV | HRV-A | 593 | HRV-A | 749 | HRV-A | 538 | HRV-A | 446 |
| 1I6 | HRV | HRV-C | 6,543 | HRV-C | 7,801 | HRV-C | 6,470 | HRV-C | 6,335 |
| 1I8 | HRV | HRV-C | 898 | HRV-C | 1,026 | HRV-C | 839 | HRV-C | 790 |
| 2A2 | HCoV 229E | HRV-C | 633 | HRV-C | 770 | HRV-C | 631 | HRV-C | 570 |
| 2A3 | HCoV 229E | HCoV 229E | 15,867 | HCoV 229E | 25,213 | HCoV 229E | 23,693 | HCoV 229E | 27,414 |
| 2A4 | HCoV 229E | HCoV 229E | 4,671 | HCoV 229E | 7,515 | HCoV 229E | 6,995 | HCoV 229E | 8,032 |
| 2A6 | HRV | HRV-C | 5,050 | HRV-C | 6,136 | HRV-C | 5,001 | HRV-C | 4,889 |
| 2A9 | hMPV | hMPV | 5,312 | hMPV | 8,428 | hMPV | 7,541 | hMPV | 8,593 |
| 2B4 | RSV | RSV | 22,783 | RSV | 40,911 | RSV | 38,309 | RSV | 47,797 |
| 2B6 | RSV | RSV | 419 | RSV | 742 | RSV | 616 | RSV | 752 |
| 2B7 | HRV | HRV-A | 3,201 | HRV-A | 4,458 | HRV-A | 3,639 | HRV-A | 3,596 |
| 2B9 | HRV | RSV | 676 | RSV | 1,105 | RSV | 910 | RSV | 1,086 |
| 2C1 | HRV | HRV-A | 5,584 | HRV-A | 7,631 | HRV-A | 6,236 | HRV-A | 6,071 |
| 2C3 | HCoV 229E | HCoV 229E | 1,915 | HCoV 229E | 3,169 | HCoV 229E | 2,944 | HCoV 229E | 3,454 |
| 2C4 | HCoV 229E | HRV-A | 228 | HRV-A | 264 | HRV-A | 229 | HRV-A | 236 |
| 2D2 | HCoV 229E | HCoV 229E | 8,883 | HCoV 229E | 14,279 | HCoV 229E | 13,440 | HCoV 229E | 15,487 |
| 2D3 | HCoV OC43 | HRV-A | 477 | HRV-A | 438 | HRV-A | 240 | HRV-A | 153 |
| 2D4 | PIV-2 | HRV-A | 562 | HRV-A | 579 | HRV-A | 336 | HRV-A | 210 |
| 2D5 | HCoV OC43 | HCoV OC43 | 16,576 | HCoV OC43 | 26,870 | HCov OC43 | 25,298 | HCoV OC43 | 29,277 |
| 2D6 | HRV | HRV-A | 3,556 | HRV-A | 3,700 | HRV-A | 2,615 | HRV-A | 2,324 |
