## Supplementary Table S4 for "DisCVR: Rapid viral diagnosis from high-throughput sequencing data"

**Table S4. Comparison between the DisCVR results and the BLAST results from the original manuscript for healthy patients and patients with unexplained acute febrile illness**

| Sample_Name | RUN | Patient status | Number of *k*-mers matching | Number of distinct k-mers matching | Top Virus hits | Number of k-mers matching the 2nd hit | 2nd Virus Hit | Top BLAST hit | Number of reads matching with BLAST | Concordance |
|---|---|---|---|---|---|---|---|---|---|---|
| 3 | SRR1748140 | healthy | NA | NA | NA | NA | NA | Human.adenovirus.C | 10 | no |
| 4 | SRR1748141 | healthy | NA | NA | NA | NA | NA | Human.adenovirus.C | 65 | no |
| 8 | SRR1748143 | healthy | 943 | 379 | Human mastadenovirus C | | | Human.adenovirus.C | 65 | yes |
| 10 | SRR1748147 | healthy | 1669 | 133 | Human immunodeficiency virus 1 | | | Human.adenovirus.C | 26 | no |
| 14 | SRR1748151 | healthy | NA | NA | NA | NA | NA | Human.adenovirus.C | 4 | no |
| 15 | SRR1748152 | healthy | NA | NA | NA | NA | NA | Human.adenovirus.C | 6 | no |
| 108 | SRR1748162 | healthy | NA | NA | NA | NA | NA | Human.adenovirus.C | 6 | no |
| 137 | SRR1748172 | healthy | 15343 | 4714 | Human immunodeficiency virus 1 | | | Human.immunodeficiency.virus | 314 | yes |
| 173 | SRR1748173 | healthy | NA | NA | NA | NA | NA | Heterosigma.akashiwo.RNA.virus | 20 | no |
| 245 | SRR1748177 | healthy | NA | NA | NA | NA | NA | Heterosigma.akashiwo.RNA.virus | 17 | no |
| 312 | SRR1748178 | healthy | 2843 | 4 | Human T-lymphotropic virus 1 | 1282 | Human mastadenovirus C | Human.adenovirus.C | 70 | no |
| 316 | SRR1748179 | healthy | 22915 | 5941 | Human immunodeficiency virus 1 | | | Human.immunodeficiency.virus | 342 | yes |
| 319 | SRR1748180 | healthy | 2247683 | 21201 | GB virus C | | | GB.virus.C | 20629 | yes |
| 325 | SRR1748181 | healthy | 1006 | 177 | Human immunodeficiency virus 1 | | | Heterosigma.akashiwo.RNA.virus | 60 | no |
| 329 | SRR1748182 | healthy | 1513 | 124 | Human immunodeficiency virus 1 | | | Human.adenovirus.C | 47 | no |
| 330 | SRR1748183 | healthy | NA | NA | NA | NA | NA | Human.adenovirus.C | 8 | no |
| 338 | SRR1748184 | healthy | 1269 | 400 | Human mastadenovirus C | | | Human.adenovirus.C | 63 | yes |
| 454 | SRR1748185 | healthy | 8570 | 4 | Human T-lymphotropic virus 1 | | | Heterosigma.akashiwo.RNA.virus | 48 | no |
| 474 | SRR1748186 | healthy | NA | NA | NA | NA | NA | Heterosigma.akashiwo.RNA.virus | 31 | no |
| 513 | SRR1748187 | healthy | NA | NA | NA | NA | NA | Heterosigma.akashiwo.RNA.virus | 38 | no |
| 582 | SRR1748189 | healthy | NA | NA | NA | NA | NA | Heterosigma.akashiwo.RNA.virus | 34 | no |
| 591 | SRR1748191 | healthy | NA | NA | NA | NA | NA | Heterosigma.akashiwo.RNA.virus | 10 | no |
| 602 | SRR1748193 | healthy | 969401 | 8234 | Hepatitis B virus | 396268 | Human immunodeficiency | Human.immunodeficiency.virus | 5408 | no |
| 630 | SRR1748196 | healthy | 11275 | 3180 | GB virus C | | | GB.virus.C | 198 | yes |
| 631 | SRR1748197 | healthy | 10230 | 226 | Human gammaherpesvirus 4 | | | Human.herpesvirus.4 | 65 | yes |
| 632 | SRR1748198 | healthy | 1242 | 324 | Human mastadenovirus C | | | Human.adenovirus.C | 68 | yes |
| 852 | SRR1748199 | healthy | 1775883 | 15597 | GB virus C | | | GB.virus.C | 16162 | yes |
| 989 | SRR1748200 | healthy | 886 | 173 | Human mastadenovirus C | | | Human.adenovirus.C | 34 | yes |
| 996 | SRR1748201 | healthy | 3359 | 300 | Human mastadenovirus C | | | Human.adenovirus.C | 199 | yes |
| 1001 | SRR1748202 | healthy | NA | NA | NA | NA | NA | Human.adenovirus.C | 18 | no |
| 1011 | SRR1748203 | healthy | NA | NA | NA | NA | NA | Human.adenovirus.C | 24 | no |
| 1014 | SRR1748204 | healthy | NA | NA | NA | NA | NA | Human.adenovirus.C | 38 | no |
| 1017 | SRR1748205 | healthy | 2712372 | 26330 | GB virus C | | | GB.virus.C | 30776 | yes |
| 1023 | SRR1748206 | healthy | 5851 | 416 | Human mastadenovirus C | | | Human.adenovirus.C | 332 | yes |
| 1032 | SRR1748207 | healthy | 122060 | 7744 | GB virus C | | | GB.virus.C | 2380 | yes |
| 1045 | SRR1748208 | healthy | NA | NA | NA | NA | NA | Agaricus.bisporus.virus.X | 14 | no |
| 1049 | SRR1748209 | healthy | 2335 | 434 | Human mastadenovirus C | | | Human.adenovirus.C | 125 | yes |
| 1051 | SRR1748210 | healthy | 1661 | 639 | XMRV-related viruses | | | Murine.leukemia.virus | 44 | yes |
| 1052 | SRR1748211 | healthy | 1984 | 345 | Human mastadenovirus C | | | Human.adenovirus.C | 140 | yes |
| 1055 | SRR1748212 | healthy | 1787 | 314 | Human mastadenovirus C | | | Human.adenovirus.C | 115 | yes |
| 1056 | SRR1748213 | healthy | 1251 | 299 | Human mastadenovirus C | | | Human.adenovirus.C | 70 | yes |
| 1059 | SRR1748214 | healthy | NA | NA | NA | NA | NA | Human.adenovirus.C | 12 | no |
| 1068 | SRR1748215 | healthy | NA | NA | NA | NA | NA | Human.adenovirus.C | 12 | no |
| 1070 | SRR1748216 | healthy | 133468 | 11227 | Human immunodeficiency virus 1 | | | Human.immunodeficiency.virus | 2446 | yes |
| 1071 | SRR1748217 | healthy | 3259 | 288 | Human mastadenovirus C | | | Human.adenovirus.C | 177 | yes |
| 1072 | SRR1748218 | healthy | 1800 | 783 | Lassa mammarenavirus | | | Lassa.virus | 40 | yes |
| 1078 | SRR1748219 | healthy | 2368 | 548 | Human mastadenovirus C | | | Human.adenovirus.C | 171 | yes |
| 1079 | SRR1748220 | healthy | 7139 | 2530 | Lassa mammarenavirus | | | Lassa.virus | 112 | yes |
| 1081 | SRR1748221 | healthy | NA | NA | NA | NA | NA | Human.adenovirus.C | 10 | no |
| 1083 | SRR1748222 | healthy | 1865 | 308 | Human mastadenovirus C | | | Human.adenovirus.C | 50 | yes |
| 1085 | SRR1748223 | healthy | 2391557 | 23939 | GB virus C | | | GB.virus.C | 32044 | yes |
| 1086 | SRR1748224 | healthy | 1779 | 390 | Human mastadenovirus C | | | Human.adenovirus.C | 80 | yes |
| 1088 | SRR1748225 | healthy | NA | NA | NA | NA | NA | Human.adenovirus.C | 12 | no |
| 1089 | SRR1748226 | healthy | NA | NA | NA | NA | NA | Cercopithecine.herpesvirus | 4 | no |
| 1093 | SRR1748227 | healthy | 214045 | 8706 | GB virus C | | | GB.virus.C | 4536 | yes |
| 1094 | SRR1748228 | healthy | 2340 | 1014 | GB virus C | | | GB.virus.C | 24 | yes |
| 1098 | SRR1748229 | healthy | 46605 | 6372 | GB virus C | | | GB.virus.C | 888 | yes |
| 1099 | SRR1748230 | healthy | NA | NA | NA | NA | NA | Human.adenovirus.C | 28 | no |
| 1102 | SRR1748231 | healthy | 1955 | 385 | Human mastadenovirus C | | | Human.adenovirus.C | 115 | yes |
| 1103 | SRR1748232 | healthy | 2633 | 512 | Human mastadenovirus C | | | Human.adenovirus.C | 198 | yes |
| 1104 | SRR1748233 | healthy | 3217447 | 21305 | GB virus C | | | GB.virus.C | 27082 | yes |
| 1106 | SRR1748234 | healthy | NA | NA | NA | NA | NA | Human.adenovirus.C | 17 | no |
| 1107 | SRR1748235 | healthy | 126054 | 9773 | Lassa mammarenavirus | | | Lassa.virus | 1350 | yes |
| 1113 | SRR1748236 | healthy | 1150 | 209 | Human mastadenovirus C | | | Human.adenovirus.C | 73 | yes |
| 1116 | SRR1748237 | healthy | 8541941 | 35544 | GB virus C | | | GB.virus.C | 94785 | yes |
| 1120 | SRR1748238 | healthy | 2433 | 404 | Human mastadenovirus C | | | Human.adenovirus.C | 118 | yes |
| 1122 | SRR1748239 | healthy | 1851 | 280 | Human mastadenovirus C | | | Human.adenovirus.C | 90 | yes |
| 1123 | SRR1748240 | healthy | 2344 | 310 | Human immunodeficiency virus 1 | | | Human.adenovirus.C | 11 | no |
| 1130 | SRR1748241 | healthy | NA | NA | NA | NA | NA | Human.adenovirus.C | 2 | no |
| 1133 | SRR1748242 | healthy | NA | NA | NA | NA | NA | Human.adenovirus.C | 16 | no |
| 1135 | SRR1748243 | healthy | 1786 | 335 | Human mastadenovirus C | | | Human.adenovirus.C | 116 | yes |
| 2000 | SRR1748244 | healthy | 1388 | 200 | XMRV-related viruses | | | Human.adenovirus.C | 56 | no |
| 2001 | SRR1748245 | healthy | 119232 | 5887 | Human immunodeficiency virus 1 | | | Human.immunodeficiency.virus | 2397 | yes |
| 2002 | SRR1748246 | healthy | NA | NA | NA | NA | NA | Human.adenovirus.C | 21 | no |
| 2003 | SRR1748247 | healthy | 908 | 132 | Human mastadenovirus C | | | Human.adenovirus.C | 36 | yes |
| 2004 | SRR1748248 | healthy | NA | NA | NA | NA | NA | Human.adenovirus.C | 23 | no |
| 2017 | SRR1748249 | healthy | NA | NA | NA | NA | NA | Human.adenovirus.C | 28 | no |
| 2018 | SRR1748250 | healthy | 1424 | 524 | GB virus C | | | GB.virus.C | 28 | yes |
| 2019 | SRR1748251 | healthy | NA | NA | NA | NA | NA | Human.adenovirus.C | 16 | no |
| 2035 | SRR1748252 | healthy | 1191926 | 14615 | GB virus C | | | GB.virus.C | 21667 | yes |
| 2036 | SRR1748253 | healthy | 910 | 92 | Human immunodeficiency virus 1 | | | Human.adenovirus.C | 53 | no |
| 2038 | SRR1748254 | healthy | 1058 | 140 | Human mastadenovirus C | | | Human.adenovirus.C | 84 | yes |
| 2039 | SRR1748255 | healthy | NA | NA | NA | NA | NA | Human.adenovirus.C | 23 | no |
| 2040 | SRR1748256 | healthy | NA | NA | NA | NA | NA | Human.adenovirus.C | 74 | no |
| 2041 | SRR1748257 | healthy | 3538 | 402 | Human mastadenovirus C | | | Human.adenovirus.C | 337 | yes |
| 2045 | SRR1748258 | healthy | NA | NA | NA | NA | NA | Human.immunodeficiency.virus | 12 | no |
| 2047 | SRR1748259 | healthy | 3084 | 365 | Human mastadenovirus C | | | Human.adenovirus.C | 217 | yes |
| 2048 | SRR1748260 | healthy | 22091 | 4172 | GB virus C | | | GB.virus.C | 424 | yes |
| 2052 | SRR1748261 | healthy | 9792 | 2441 | Lassa mammarenavirus | | | Lassa.virus | 166 | yes |
| 2070 | SRR1748262 | healthy | 17718 | 5009 | Lassa mammarenavirus | | | Lassa.virus | 252 | yes |
| 102C | SRR1748159 | afebrile | NA | NA | NA | NA | NA | Human.adenovirus.C | 4 | no |
| 102F | SRR1748160 | afebrile | NA | NA | NA | NA | NA | Human.adenovirus.C | 2 | no |
| 102M | SRR1748161 | afebrile | NA | NA | NA | NA | NA | Human.adenovirus.C | 12 | no |
| 111C | SRR1748163 | afebrile | 2782 | 131 | Human immunodeficiency virus 1 | | | Human.adenovirus.C | 12 | no |
| 111F | SRR1748164 | afebrile | 5962 | 232 | Human immunodeficiency virus 1 | 1481 | Human mastadenovirus C | Human.adenovirus.C | 55 | no |
| 111M | SRR1748165 | afebrile | 1019 | 189 | Human mastadenovirus C | | | Human.adenovirus.C | 50 | yes |
| 114C | SRR1748166 | afebrile | NA | NA | NA | NA | NA | Human.adenovirus.C | 67 | no |
| 114F | SRR1748167 | afebrile | 1023 | 149 | Human immunodeficiency virus 1 | | | Human.adenovirus.C | 22 | no |
| 114M | SRR1748168 | afebrile | 1243 | 180 | Human immunodeficiency virus 1 | | | Human.adenovirus.C | 21 | no |
| 11C | SRR1748148 | afebrile | 18072 | 1382 | Human mastadenovirus C | | | Human.adenovirus.C | 1083 | yes |
| 11F | SRR1748149 | afebrile | 17069 | 1392 | Human mastadenovirus C | | | Human.adenovirus.C | 986 | yes |
| 11M | SRR1748150 | afebrile | 18473 | 1367 | Human mastadenovirus C | | | Human.adenovirus.C | 1079 | yes |
| 120C | SRR1748169 | afebrile | NA | NA | NA | NA | NA | Human.adenovirus.C | 26 | no |
| 120F | SRR1748170 | afebrile | NA | NA | NA | NA | NA | Human.adenovirus.C | 23 | no |
| 120M | SRR1748171 | afebrile | NA | NA | NA | NA | NA | Human.adenovirus.C | 16 | no |
| 1C | SRR1748134 | afebrile | NA | NA | NA | NA | NA | Human.adenovirus.C | 40 | no |
| 1F | SRR1748135 | afebrile | 3785 | 720 | Human mastadenovirus C | | | Human.adenovirus.C | 226 | yes |
| 1M | SRR1748136 | afebrile | 3466 | 585 | Human mastadenovirus C | | | Human.adenovirus.C | 159 | yes |
| 230C | SRR1748174 | afebrile | 885 | 170 | Human mastadenovirus C | | | Human.adenovirus.C | 41 | yes |
| 230F | SRR1748175 | afebrile | 1586 | 168 | Human immunodeficiency virus 1 | 1199 | Human mastadenovirus C | Human.adenovirus.C | 66 | no |
| 230M | SRR1748176 | afebrile | 4337 | 245 | Human immunodeficiency virus 1 | | | Human.adenovirus.C | 44 | no |
| 2C | SRR1748137 | afebrile | 3489 | 566 | Human mastadenovirus C | | | Human.adenovirus.C | 168 | yes |
| 2F | SRR1748138 | afebrile | 8763 | 299 | Human immunodeficiency virus 1 | 1058 | Human mastadenovirus C | Human.adenovirus.C | 60 | no |
| 2M | SRR1748139 | afebrile | 5100 | 172 | Human immunodeficiency virus 1 | 3081 | Human mastadenovirus C | Human.adenovirus.C | 145 | no |
| 49C_HiSeq | SRR1748153 | afebrile | 56859 | 275 | Human immunodeficiency virus 1 | 20838 | Human mastadenovirus C | Human.adenovirus.C | 444 | no |
| 49F | SRR1748154 | afebrile | 3909 | 977 | Human mastadenovirus C | | | Human.adenovirus.C | 228 | yes |
| 49M | SRR1748155 | afebrile | 4250 | 1111 | Human mastadenovirus C | | | Human.adenovirus.C | 271 | yes |
| 54C | SRR1748156 | afebrile | NA | NA | NA | NA | NA | Human.adenovirus.C | 8 | no |
| 54F | SRR1748157 | afebrile | NA | NA | NA | NA | NA | GB.virus.C | 2 | no |
| 54M | SRR1748158 | afebrile | NA | NA | NA | NA | NA | Cafeteria.roenbergensis.virus.BV.PW | 4 | no |
| 572PP | SRR1748188 | healthy | NA | NA | NA | NA | NA | Human.adenovirus.C | 4 | no |
| 587-hex | SRR1748190 | healthy | 2297 | 445 | Human mastadenovirus C | | | Human.adenovirus.C | 99 | yes |
| 592-hex | SRR1748192 | healthy | 2217 | 468 | Human mastadenovirus C | | | Human.adenovirus.C | 124 | yes |
| 5Healthy_243M_Pi | SRR1748142 | afebrile | 1099740 | 18396 | GB virus C | | | GB.virus.C | 11403 | yes |
| 620PP | SRR1748195 | healthy | NA | NA | NA | NA | NA | Human.adenovirus.C | 40 | no |
| 9C | SRR1748144 | afebrile | 23574 | 1402 | Human mastadenovirus C | | | Human.adenovirus.C | 1335 | yes |
| 9F | SRR1748145 | afebrile | 20314 | 1408 | Human mastadenovirus C | | | Human.adenovirus.C | 1334 | yes |
| 9M | SRR1748146 | afebrile | 13309 | 1357 | Human mastadenovirus C | | | Human.adenovirus.C | 752 | yes |
| DF_LIB1 | SRR1748263 | afebrile | NA | NA | NA | NA | NA | Human.adenovirus.C | 15 | no |
| DF_LIB2 | SRR1748264 | afebrile | 17284365 | 68280 | GB virus C | | | GB.virus.C | 138614 | yes |
| DF_LIB3 | SRR1748265 | afebrile | 9263007 | 48169 | GB virus C | | | GB.virus.C | 66847 | yes |
| DF_LIB4 | SRR1748266 | afebrile | NA | NA | NA | NA | NA | Human.adenovirus.C | 6 | no |
| DF_LIB5 | SRR1748267 | afebrile | 3318817 | 38170 | GB virus C | | | GB.virus.C | 31146 | yes |
| DF_LIB6 | SRR1748268 | afebrile | 709533 | 10537 | GB virus C | | | GB.virus.C | 6825 | yes |
| DF_LIB7 | SRR1748269 | afebrile | 60760 | 8548 | Human immunodeficiency virus 1 | | | Human.immunodeficiency.virus | 815 | yes |
| DF_LIB8 | SRR1748270 | afebrile | 4364508 | 27905 | GB virus C | | | GB.virus.C | 31573 | yes |
| DFultra1 | SRR1748271 | afebrile | NA | NA | NA | NA | NA | Human.adenovirus.C | 18 | no |
| FUOP1 | SRR1748273 | healthy | 832705 | 11465 | GB virus C | | | GB.virus.C | 8676 | yes |
| FUOP1_LIB11-18 | SRR1748272 | healthy | 3458159 | 29008 | GB virus C | | | GB.virus.C | 63229 | yes |
| FUOP2 | SRR1748275 | healthy | 132512 | 8400 | GB virus C | | | GB.virus.C | 1778 | yes |
| FUOP2_LIB11-18 | SRR1748274 | healthy | 1047 | 167 | Human immunodeficiency virus 1 | | | Human.adenovirus.C | 62 | no |
| FUOP3 | SRR1748277 | healthy | NA | NA | NA | NA | NA | Human.adenovirus.C | 18 | no |
| FUOP3_LIB11-18 | SRR1748276 | healthy | 2153 | 132 | Human immunodeficiency virus 1 | 1514 | Human mastadenovirus C | Human.adenovirus.C | 55 | no |
| FUOP4 | SRR1748279 | healthy | 29068 | 5811 | Lassa mammarenavirus | | | Lassa.virus | 430 | yes |

| | | | | | | | | | | |
|---|---|---|---|---|---|---|---|---|---|---|
| FUOP4_LIB11-18 | SRR1748278 | healthy | 196059 | 7705 | GB virus C | | | GB.virus.C | 5014 | yes |
| FUOP5 | SRR1748280 | healthy | NA | NA | NA | NA | NA | Human.adenovirus.C | 10 | no |
| FUOP6 | SRR1748281 | healthy | NA | NA | NA | NA | NA | Human.adenovirus.C | 40 | no |
| FUOP7 | SRR1748282 | healthy | 931111 | 12880 | GB virus C | | | GB.virus.C | 18230 | yes |
| FUOP8 | SRR1748283 | healthy | 21956 | 5104 | GB virus C | | | GB.virus.C | 380 | yes |
| H_ultra_p1 | SRR1748284 | afebrile | 1105645 | 22112 | GB virus C | | | GB.virus.C | 18224 | yes |
| H_ultra_p2 | SRR1748285 | afebrile | 1271189 | 21875 | GB virus C | | | GB.virus.C | 19352 | yes |
| HP1 | SRR1748287 | afebrile | 354927 | 8497 | GB virus C | | | GB.virus.C | 5866 | yes |
| HP1_LIB11-18 | SRR1748286 | afebrile | 28025 | 4130 | GB virus C | | | GB.virus.C | 516 | yes |
| HP2 | SRR1748289 | afebrile | 61919 | 6142 | GB virus C | | | GB.virus.C | 952 | yes |
| HP2_LIB11-18 | SRR1748288 | afebrile | 16066 | 3540 | GB virus C | | | GB.virus.C | 384 | yes |
| HP3 | SRR1748290 | afebrile | 2006 | 613 | Human mastadenovirus C | | | Human.adenovirus.C | 158 | yes |
| HP4 | SRR1748291 | afebrile | 8019 | 975 | Human mastadenovirus C | | | Human.adenovirus.C | 445 | yes |
| HP5 | SRR1748292 | afebrile | 2018 | 563 | Human mastadenovirus C | | | Human.adenovirus.C | 118 | yes |
| HP6 | SRR1748293 | afebrile | 4565 | 698 | Human mastadenovirus C | | | Human.adenovirus.C | 327 | yes |
| HP7 | SRR1748294 | afebrile | 1122059 | 14686 | GB virus C | | | GB.virus.C | 20164 | yes |
| HP8 | SRR1748295 | afebrile | 2258 | 718 | Human mastadenovirus C | | | Human.adenovirus.C | 130 | yes |
| LIB10_pool1 | SRR1748296 | afebrile | 11883 | 1351 | Human mastadenovirus C | | | Human.adenovirus.C | 785 | yes |
| LIB10_pool10 | SRR1748305 | healthy | 18494 | 4600 | Human immunodeficiency virus 1 | | | Human.immunodeficiency.virus | 305 | yes |
| LIB10_pool11 | SRR1748306 | healthy | NA | NA | NA | NA | NA | Human.adenovirus.C | 49 | no |
| LIB10_pool12 | SRR1748307 | healthy | 2237 | 762 | Human mastadenovirus C | | | Human.adenovirus.C | 157 | yes |
| LIB10_pool13 | SRR1748308 | healthy | 1202 | 412 | Human mastadenovirus C | | | Human.adenovirus.C | 79 | yes |
| LIB10_pool14 | SRR1748309 | healthy | 173928 | 6771 | GB virus C | | | GB.virus.C | 2193 | yes |
| LIB10_pool15 | SRR1748310 | healthy | NA | NA | NA | NA | NA | Acyrthosiphon.pisum.virus | 0 | no |
| LIB10_pool16 | SRR1748311 | afebrile | 56805 | 275 | Human immunodeficiency virus 1 | 17509 | Human.adenovirus.C | Human.adenovirus.C | 1129 | no |
| LIB10_pool2 | SRR1748297 | healthy | 26163 | 5854 | Human immunodeficiency virus 1 | | | Human.immunodeficiency.virus | 516 | yes |
| LIB10_pool3 | SRR1748298 | healthy | 968203 | 15270 | GB virus C | | | GB.virus.C | 10271 | yes |
| LIB10_pool4 | SRR1748299 | healthy | 6654 | 1355 | Human mastadenovirus C | | | Human.adenovirus.C | 384 | yes |
| LIB10_pool5 | SRR1748300 | healthy | 29689 | 5233 | GB virus C | | | GB.virus.C | 424 | yes |
| LIB10_pool6 | SRR1748301 | healthy | 13370 | 1744 | Hepatitis C virus | | | Hepatitis.C.virus | 1208 | yes |
| LIB10_pool7 | SRR1748302 | healthy | 4904 | 1119 | Human mastadenovirus C | | | Human.adenovirus.C | 311 | yes |
| LIB10_pool8 | SRR1748303 | healthy | NA | NA | NA | NA | NA | Human.adenovirus.C | 19 | no |
| LIB10_pool9 | SRR1748304 | healthy | NA | NA | NA | NA | NA | Human.adenovirus.C | 20 | no |
